# Towards understanding biology of leydiogioma. G protein-coupled receptor and peroxisome proliferator-activated receptor crosstalk regulates lipid metabolism and steroidogenesis in Leydig cell tumors

**DOI:** 10.1101/477901

**Authors:** Malgorzata Kotula-Balak, Ewelina Gorowska-Wojtowicz, Agnieszka Milon, Piotr Pawlicki, Alicja Kaminska, Laura Pardyak, Waclaw Tworzydlo, Bartosz J. Płachno, Anna Hejmej, Jan K. Wolski

## Abstract

Leydig cell tumors (LCT) are the most common type of testicular sex cord-stromal tumor. In this report, we implicate the G-coupled estrogen receptor (GPER) and peroxisome proliferator receptor (PPAR) in regulation of lipid homeostasis and the expression of steroidogenesis-controlling molecules in clinical specimens of LCTs and cell line (mouse tumor Leydig cells; MA-10). We also show the general structure and morphology of human LCTs with the use of scanning electron microscopy and light microscopy, respectively. In LCTs, protein immunoblotting and immunohistochemical analysis revealed increased expression of GPER and decreased expression of PPARα, β and γ. Concomitantly, changes in expression pattern of the lutropin receptor (LHR), protein kinase A (PKA), perilipin (PLIN), hormone sensitive lipase (HSL), steroidogenic acute regulatory protein (StAR), translocator protein (TSPO), HMG-CoA synthase (HMGCA), and HMG-CoA reductase (HMGCR) were observed.

Using MA-10 cells treated with GPER and PPAR antagonists (alone and in combination), we demonstrated there is a GPER-PPAR mediated control of cholesterol concentration. In addition, GPER-PPARα regulated estradiol secretion, while GPER-PPARγ affected cGMP concentration. It is assumed that GPER and PPAR can be altered in LCT, resulting in a perturbed lipid balance and steroidogenesis. In LCTs, the phosphatidylinositol-3-kinase (PI3K)-Akt-mTOR signaling pathway was disturbed. Thus, PI3K-Akt-mTOR, together with cGMP, can play a role in LCT proliferation, growth, and metastasis as well as lipid balance control.

In conclusion, we discuss the implications of GPER-PPAR interaction with lipid metabolism and steroidogenesis controlling-molecules in LCT biology that can be used in future studies as potential targets of diagnostic and therapeutic implementations.

## Introduction

Leydig cell tumor (LCT; leydigioma) is the most common non-germ cell gonadal tumor, accounting for 1-3% of all testicular tumors in adults and 4-9% in prepubertal children [1-3]. In recent years, a marked increase in the incidence of LCTs has been observed (14.7% of all testicular tumors removed). LCTs are usually benign tumors especially in infancy [4, 5], although local recurrence or metachronous tumors of the contralateral testis have also been described. Survival after diagnosis of primary LCTs ranged from 2 months to 17 years [6]. Patients with LCTs usually have symptoms of testicular swelling [7] or various endocrinological disruptions [8]. Some patients have associated issues with endocrine symptoms e.g. gynecomastia and decreased libido. Gynecomastia is the main clinical manifestation in adults, but it may also be clinically significant in affected children who undergo precocious puberty [9]. In the latter, behavioral perturbations were observed [10]. Some cases of LCTs were linked with increased plasma estradiol concentrations and were merely revealed by gynecomastia [11,12]. Moreover, infertility and azoospermia are not an unusual finding in patients with LCTs [13].

Less than 0.2% of all testicular cancers were evidenced by metastatic spread. Besides retroperitoneal nodes, other metastatic sites are liver, lungs, bone, and mediastinum [14, 15]. In prepubertal patients, even malignant LCTs are less likely to metastasize. Radical orchiectomy is the current standard of care but testis sparing surgery (TSS) (enucleation), in conjunction with intraoperative frozen section (FSE), has been recently attempted with promising results [16]. Prepubertal individuals and those with smaller tumors that lack evidence of malignancy are directly recommended for TSS.

The etiology of LCTs is unknown and appears heterogeneous. Furthermore, the molecular basis of LCTs is poorly understood. Some studies showed a possible role of genetic factors in LCT development. Interestingly, genetic mutations identified in children and adults were different and, in some cases, associated with other cancers [17]. In adults, it was observed that a somatic activating mutation in the guanine nucleotide-binding protein α gene may result in tumor development, leading to overexpression of the inhibin α subunit and hyperactivity of sex steroid biosynthesis [5]. In addition, Carvajal-Carmona *et al.*, [18] reported an inherited fumarate hydratase mutation appears to cause tumor growth through activation of the hypoxia pathway. According to study of Lejeune *et al.*, [11] alterations in local stimuli, including Müllerian-inhibiting hormone, inhibin, growth factors, and temperature, may also create favorable conditions for initiation and development of LCTs.

Decreased Leydig cell function is common in men with reproductive disorders, including testicular dysgenesis syndrome (TDS). This syndrome is comprised of subfertility, cryptorchidism, hypospadias, and testicular cancer that is pathogenetically linked to impaired testis development and function [19, 20]. Leydig cell impairment manifests as a decreased testosterone/lutropin (LH) ratio and the presence of Leydig cell micronodules in the testis [21]. Additionally, uncontrolled synthesis and secretion of testosterone may suppress LH secretion and impair spermatogenesis [22]. The number and size of micronodules (clusters-when Leydig cell number >15) increases with the severity of TDS and increasing gonadotrophin levels [21, 23]. Due to ultrastructural changes demonstrated in Leydig cells within micronodules (decreased smooth endoplasmic reticulum, irregularly indented nuclear membrane, decreased lipofuscin pigment granules, and Reinke crystals) failure of their maturation is suggested [24, 25]. Also, the proportion of morphologically abnormal Leydig cells was inversely correlated with testosterone levels in patients with primary testicular disorders such as cryptorchidism, Klinefelter syndrome, and Del Castillo's syndrome [26].

Besides micronodules, other histopathological changes of Leydig cells have been described. In patients with germ cell tumors, Leydig cell hypertrophy and hyperplasia were linked to elevated levels of chorionic gonadotropin [27]. In addition, various chemicals induce Leydig cell hyperplasia because of disruption of the hypothalamic-pituitary-axis [28]. Neoplastic proliferation of Leydig cells may result in the synthesis of non-functional steroid hormones. Alternatively, an excess of various hormones (*e.g*. estrogen, prolactin) produce elevated LH levels that excessively stimulate steroidogenic Leydig cell function [29]. Concomitantly, in experimental studies of animals exposed to hormones, hormonally active chemicals, or other hormonal modulators, Leydig cell hypertrophy was demonstrated [30-33]. However, there is no evidence as to whether Leydig cell pathology including micronodules, hyperplasia, and/or hypertrophy may further develop into leydigioma [34].

Mitogenicity associated with estrogen receptor-mediated cellular events is believed to be the mechanism by which estrogens contribute to tumorigenesis. Currently, implications of estrogen signaling *via* canonical estrogen receptors (ERs), G-protein coupled membrane estrogen receptor (GPER), as well as estrogen-related receptors (ERRs) is recognized in animal and human Leydig cell tumorigenesis [35-37]. Until recently, the function of GPER in testicular cells was only partially known. In human testis, Fietz *et al*. [38, 39] showed high levels of GPER mRNA expression in Leydig cells. In fact, not only is GPER able to bind sex steroids but also binds the peroxisome proliferator-activated receptors (PPARs) [40]. In amphibians, rodents, and humans, three forms of PPAR have been described to date: PPARα, PPARβ (also known as PPARδ), and PPARγ [41]. PPARs target genes that encode enzymes involved in peroxisome and mitochondria function as well as those of fatty acids, apolipoproteins, and lipoprotein lipase. Little is known about PPARs in the male reproductive system. In rat testis, PPARs are mainly expressed in Leydig and Sertoli cells [42]. It was shown that some PPAR chemicals alter testosterone production [43], and their long-term administration results in Leydig cell tumor development in rats [44, 45].

Scarce data are available on the molecular and biochemical characteristics of LCTs. Maintaining an adequate hormonal balance within the testis is the basis for proper gonadal function, thus playing a pivotal role for blocking hormone-secreting Leydig cell tumor development [46, 47]. Of note, in tumor cells, proliferation, growth, apoptosis, and communication are not only disturbed but other processes as well e.g. lipid metabolism [48].

It is worth noting that biosynthesis of sex steroids is multi-level, controlled process [49]. It requires the coordinated expression of number of genes, proteins of various function (receptors e.g. lutropin receptor; LHR, enzymes, transporters e.g. translocator protein; TSPO, steroidogenic acute regulatory protein; StAR, and regulators), signaling molecules (e.g. protein kinase A; PKA), and their regulators in response to LH stimulation. Moreover, for cellular steroidogenic function, global lipid homeostasis is crucial. Perilipin (PLIN), hormone sensitive lipase (HSL), and HMG-CoA synthase (HMGCS) and reductase (HMGCR) are members of a cell structural and enzymatic protein machinery controlling lipid homeostasis [50]. Activation of lipid metabolism is an early event in tumorigenesis, however, the precise expression pattern of lipid balance-controlling molecules and their molecular mechanism remains poorly characterized.

This study aims to determine the potential link between GPER and PPAR and whether this interaction regulates lipid homeostasis in LCTs. To further investigate the relationship of Leydig cell tumorigenesis to these receptors while elucidating the effects of their interactions mouse tumor Leydig cells (MA-10) were utilized.

## Materials and Methods

### 2.1. Tissue samples and ethical considerations

Adult samples were residual tissues from testicular biopsy (microdissection TESE, by Schlegel, 1998) specimens collected at the nOvum Fertility Clinic, Warsaw, Poland from patients (31-45 year-old; n=24) diagnosed due to azoospermia (micronodules LCTs were recognized during surgery) after written informed consent according to the approval regulations by the National Commission of Bioethics at the Jagiellonian University in Krakow, Poland; permit no. 1072.6120.218.2017 and were carried out in accordance with the Declaration of Helsinki.

After evaluation by pathologists, the remaining tissue fragments were snap-frozen or fixed and paraffin-embedded, and stored at the Department of Endocrinology, Institute of Zoology and Biomedical Research, Jagiellonian University in Krakow, Poland.

### 2.2. Cell culture and treatments

The mouse Leydig cell line MA-10 was a generous gift from Dr. Mario Ascoli (University of Iowa, Iowa City, USA), and was maintained under standard technique [51]. Middle passages (p25-p28) of MA-10 cells were used for the study. The cells were grown in Waymouth’s media (Gibco, Grand Island, NY) supplemented with 12% horse serum and 50 mg/l of gentamicin at 37 °C in 5% CO_2_. Cells were plated overnight at a density of 1 × 10^5^ cells/ml per well. Morphological and biochemical properties of MA-10 cells are regularly checked by microscopic observation, analysis of proliferation (TC20 Bio-Rad automated cell counter), mycoplasma detection (MycoFluor™ Mycoplasma Detection Kit; ThermoFisher Scientific), qRT-PCR analysis of characteristic genes and ELISA measurements of secretion products according to cell line authentication recommendations of the Global Bioresource Center (ATCC).

Twenty-four hours before the experiments, the medium was removed and replaced with a medium without phenol red supplemented with 5% dextran-coated, charcoal-treated FBS (5% DC-FBS) to exclude estrogenic effects caused by the medium. Next, cells were treated with selective antagonists: GPER [(3a*S*\*,4*R*\*,9b*R*\*)-4-(6-Bromo-1,3-benzodioxol-5-yl)-3a,4,5,9b-3*H*-cyclopenta[*c*]quinolone; G-15] (Tocris Bioscience, Bristol, UK), PPARα [*N*-((2*S*)-2-(((1*Z*)-1-Methyl-3-oxo-3-(4-(trifluoromethyl)phenyl)prop-1-enyl)amino)-3-(4-(2-(5-methyl-2-phenyl-1,3-oxazol-4-yl)ethoxy)phenyl)propyl) propanamide, GW6471] or PPARγ [2-Chloro-5-nitro-*N*-4-pyridinylbenzamide, T0070907] freshly prepared as 100nM stock solutions in dimethyl sulfoxide (DMSO) (Sigma-Aldrich) and stored at −20°C. A stock concentrations were subsequently dissolved in Waymouth’s media to a final concentrations. Cells were treated with G-15, PPARα or PPARγ alone or together for 24hours. Doses of G-15 (10nM), PPARα (10µM) or PPARγ (10µM) [52]. Control cells were treated with DMSO (final conc. 0.1%). Cell lysates and culture media were frozen in −20°C for further analyses.

### 2.3. Scanning electron microscopy analysis

LCTs were fixed in a mixture of 2.5% glutaraldehyde with 2.5% formaldehyde in a 0.05 M cacodylate buffer (Sigma; pH 7.2) for seven days, washed three times in a 0.1 M sodium cacodylate buffer and later dehydrated and subjected to critical-point drying. They were then sputter-coated with gold and examined at an accelerating voltage of 20 kV or 10 KV using a Hitachi S-4700 scanning electron microscope (Hitachi, Tokyo, Japan), which is housed in the Institute of Geological Sciences, Jagiellonian University in Kraków).

### 2.4. Histology

For routine histology, hematoxylin-eosin staining was performed on 4% paraformaldehyde LCT sections. As a control paraffin sections of human testis (cat. No. HP-401; Zyagen, San Diego, CA, USA) were used.

### 2.5. RNA isolation, reverse transcription and real-time quantitative RT-PCR

Total RNA was extracted from LCTs specimens and human Leydig cells (cat. no 10HU- 103; ixCells Biotechnologies, San Diego CA, USA) using TRIzol^®^ reagent (Life Technologies, Gaithersburg, MD, USA) according to the manufacturer’s instructions. The yield and quality of the RNA were assessed using a NanoDrop ND2000 Spectrophotometer (Thermo Scientific, Wilmington, DE, USA). Samples with a 260/280 ratio of 1.95 or greater and a 260/230 ratio of 2.0 or greater were used for analysis. Total cDNA was prepared using High-Capacity cDNA Reverse Transcription Kit (Applied Biosystems, Carlsbad, CA, USA) according to the manufacturer’s instructions.

The purified total RNA was used to generate total cDNA. A volume equivalent to 1 μg of total RNA was reverse transcribed using the High-Capacity cDNA Reverse Transcription Kit (Applied Biosystems, Carlsbad, CA, USA) according to the manufacturer's instructions. Total cDNA was prepared in a 20-μL volume using a random primer, dNTP mix, RNase inhibitor and reverse transcriptase (RT). Parallel reactions for each RNA sample were run in the absence of RT to assess genomic DNA contamination. RNase-free water was added in place of the RT product.

Real-time RT-PCR was performed using the StepOne Real-Time PCR system (Applied Biosystems) and optimized standard conditions as described previously by Kotula-Balak et al. [53, 54]. Based on the gene sequences in Ensembl database primer sets were designed using Primer3 software (Table, supplementary material). Selected primers were synthesized by Institute of Biochemistry and Biophysics, Polish Academy of Science (Warsaw, Poland). To calculate the amplification efficiency serial cDNA dilution curves were produced for all genes. A graph of threshold cycle (Ct) versus log10 relative copy number of the sample from a dilution series was produced. The slope of the curve was used to determine the amplification efficiency: %E = (10−^1/slope^-1) × 100. All PCR assays displayed efficiency between 94% and 104%.

Detection of amplification gene products was performed with 10 ng cDNA, 0.5 μM primers, and SYBR Green master mix (Applied Biosystems) in a final volume of 20 μL. Amplifications were performed as follows: 55 °C for 2 min, 94 °C for 10 min, followed by annealing temperature for 30 s (Table 1) and 45 s 72 °C to determine the cycle threshold (Ct) for quantitative measurement. To confirm amplification specificity, the PCR products from each primer pair were subjected to melting curve analysis and subsequent agarose gel electrophoresis (not shown). In all real-time RT-PCR reactions, a negative control corresponding to RT reaction without the reverse transcriptase enzyme and a blank sample were carried out. All PCR products stained with Midori Green Stain (Nippon Genetics Europe GmbH, Düren, Germany) were run on agarose gels. Images were captured using a Bio–Rad Gel Doc XR System (Bio–Rad Laboratories, Hercules, CA, USA) (not shown). mRNA expressions were normalized to the *β-actin* mRNA (relative quantification, RQ = 1) with the use of the 2^−ΔΔCt^ method.

### 2.6. Western blotting

For quantification of protein expression (Table 1) from LCTs proteins (as a control human Leydig cells; cat. No 10HU-103; ixCells Biotechnologies, San Diego CA, USA) were extracted in 50 μl of radioimmunoprecipitation assay buffer (RIPA; Thermo Scientific, Inc. Rockford IL, USA) and protease inhibitor cocktail (Sigma Chemical Co., St. Louis, Missouri, USA). Concentration of proteins was determined with Bradford reagent (Bio-Rad Protein Assay; Bio-Rad Laboratories GmbH, Munchen, Germany), using bovine serum albumin as a standard. Aliquots (50 μg protein) of cell lysates were used for electrophoresis on 12% mini gel by standard SDS-PAGE procedures and electrotransferred to polyvinylidene difluoride (PVDF) membranes (Millipore Corporate, MA, USA) by a semi-dry transfer cell (Bio-Rad). Then, blots were blocked with 5% nonfat dry milk in TBS, 0.1% Tween 20, overnight at 4 °C with shaking, followed by an incubation with respective antibodies (Table 1). The membranes were washed and incubated with a secondary antibody conjugated with the horseradish-peroxidase labeled goat anti-mouse or goat anti-rabbit IgGs (Vector Labs., Burlingame, CA, USA) at a dilution 1:1000, for 1 h at RT. Immunoreactive proteins were detected by chemiluminescence with Western Blotting Luminol Reagent (Santa Cruz Biotechnology), and images were captured with a ChemiDoc XRS + System (Bio-Rad Laboratories). All immunoblots were stripped with stripping buffer containing 62.5 mM Tris–HCl, 100 mM 2-mercaptoethanol, and 2% SDS (pH 6.7) at 50 °C for 30 min and incubated in a rabbit polyclonal antibody against β-actin. Each data point was normalized against its corresponding β-actin data point.

To obtain quantitative results, immunoblots were scanned with Image Lab 2.0 (Bio-Rad Laboratories). Then, a bound antibody was revealed using DAB as the substrate. Finally, the membranes were dried and then scanned using Epson Perfection Photo Scanner (Epson Corporation, CA, USA). Molecular masses were estimated by reference to standard proteins (Prestained SDS-PAGE Standards, Bio-Rad Labs, GmbH, Munchen, Germany). Quantitative analysis was performed for three separately repeated experiments using a public domain ImageJ software (National Institutes of Health, Bethesda, MD) as described elsewhere [55]. The relative protein levels were expressed as arbitrary units.

2.7 Immunohistochemistry

To optimize immunohistochemical staining testis sections both control (Zyagen, San Diego, CA, USA) and LCTs sections were immersed in 10 mM citrate buffer (pH 6.0) and heated in a microwave oven (2 × 5 min, 700 W). Thereafter, sections were immersed sequentially in H_2_O_2_ (3 %; v/v) for 10 min and normal goat serum (5 %; v/v) for 30 min which were used as blocking solutions. After overnight incubation at 4 °C with primary antibodies listed in Table 1. Next respective biotinylated antibodies (anti-rabbit and anti-mouse IgGs; 1: 400; Vector, Burlingame CA, USA) and avidin-biotinylated horseradish peroxidase complex (ABC/HRP; 1:100; Dako, Glostrup, Denmark) were applied in succession. Bound antibody was visualized with 3,3’-diaminobenzidine (DAB) (0.05 %; v/v; Sigma-Aldrich) as a chromogenic substrate. Control sections included omission of primary antibody and substitution by irrelevant IgG.

### 2.8. Cholesterol assay

The Amplex^®^ Red Cholesterol Assay Kit (Molecular Probes Inc., Eugene, OR, USA) was used for cholesterol content (µM) analysis in control and treated with GPER and PPAR (alone or in combinations) MA-10 cells. For measurement 100 µl cell lysates was used according to manufacturer’ s protocol. Data were expressed as mean ± SD. The fluorescence (λ = 580 nm) was measured with the use of a fluorescence multiwell plate reader SPARK Tecan, Switzerland.

### 2.9. cGMP concentration and estradiol secretion

The production of cGMP in control and treated with GPER and PPAR (alone or in combinations) MA-10 cells was measured by General Cyclic guanosine monophosphate Elisa kit assay (EIAab Wuhan Eiaab Science Co., LTD, Wuhan, China) according to the manufacturer’s instructions with detection level 0.31 to 20.0 ng/mL. The cGMP levels were calculated as ng/mL.

Estradiol Enzyme Immunoassay Kit (DRG, Inc. Int. Springfield, USA) was used for measurement of estradiol content in culture medium from control and treated with GPER and PPAR (alone or in combinations) MA-10 cells according to the manufacturer's instructions. The sensitivity of the assay was 10.6 pg/mL. The absorbance (λ = 450 nm) was measured. Data were expressed as mean ± SD.

The measurements were performed with the use ELISA apparatus (Labtech LT-4500).

### 2.10. Statistics

Three biological repeats of each sample (n = 7) and three independent experiments were performed. Each variable was tested using the Shapiro-Wilk W-test for normality. The homogeneity of variance was assessed with Levene’s test. Comparisons were performed by one-way ANOVA, followed by Dunnett’s post hoc test (GB-STAT software, v. 7.0; Dynamic Microsystems) to determine the significant differences between proteins expression levels, and cholesterol content, cGMP content and estradiol secretion. Statistical analyses were performed on raw data using Statistica 10 software (StatSoft Inc., Tulsa, OK, USA). Data were presented as means *±* S.D. Data were considered statistically significant at ∗ *p* < 0.05, ∗∗ *p* < 0.01 and ∗∗∗*p* < 0.001.

## Results

### 3.1. Scanning electron microscopic and morphological observations of LCTs

Scanning electron microscopy analyses of LCT biopsy fragments revealed the tumors are relatively compact structures of oval or slightly elongated shape (Fig.1a A-C) with tumor cells apposed and adhering to one another (Fig.1a D, E). It is important to note that some tumor cells were fused. Our observations also revealed that compact areas of tumor cells are separated by deep grooves. Between those grooves, compact tumor sheets are formed (Fig.1b E-G). Cells within sheets are tightly linked by thick projections and masses of such connections were observed between cells from neighboring tumor sheets (Fig.1b E –F). Higher magnification revealed the presence of elongated, delicate filiform fibrillar projections that form a cage-like structure covering individual tumor sheets (Fig.1b I – L).

**Figure 1.**
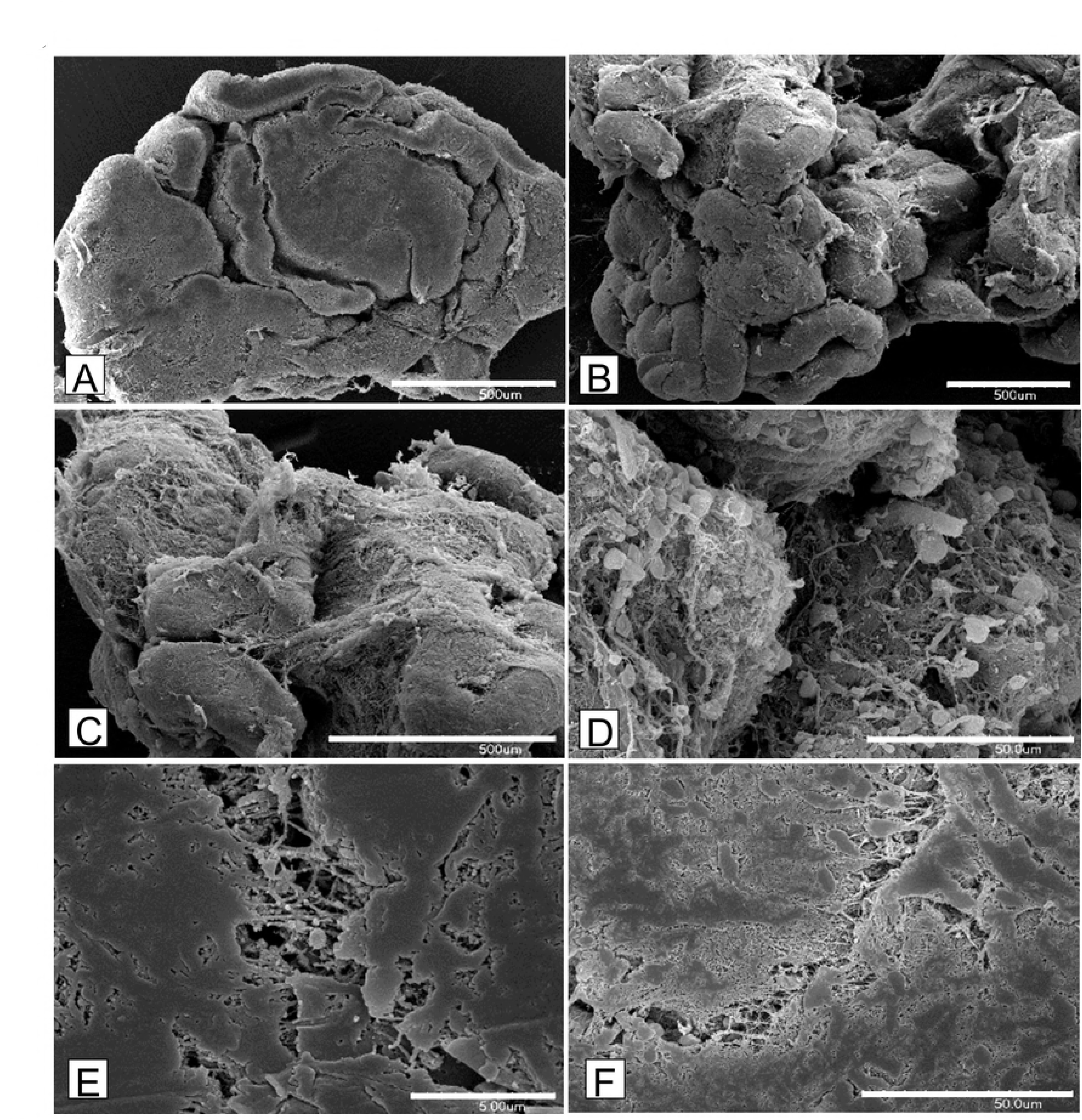
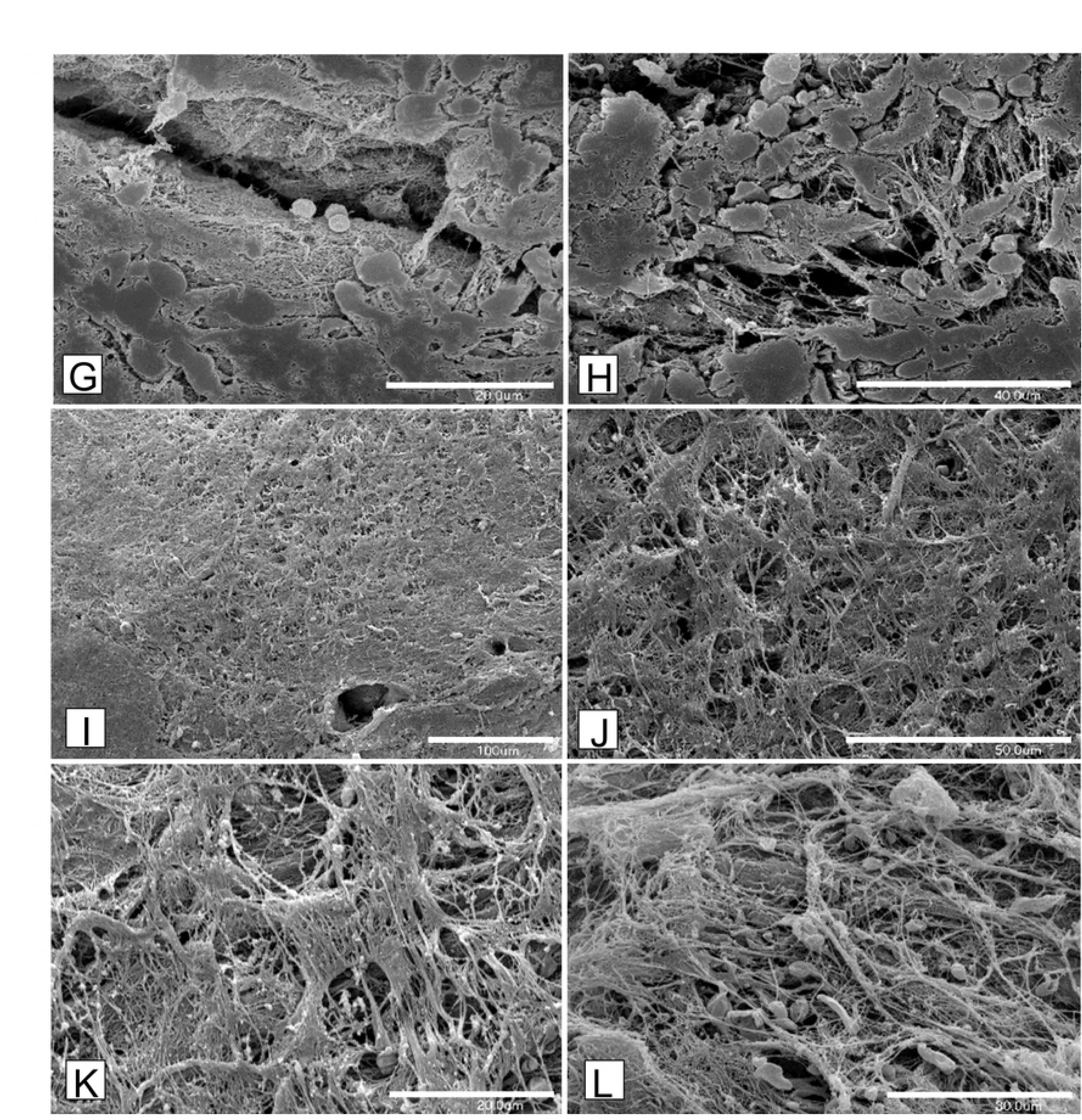
Morphology of human Leydig cell tumors – scanning electron microscopic analysis. (1a A-F and 1b G-L) Representative microphotographs of scanning electron microscopic analysis of human Leydig cell tumors (LCTs). Bars represent 1µm. For analysis n = 12 specimens were used.

Hematoxylin-eosin staining demonstrated a mixture of four types of cells in the tumor mass when compared to controls, where single or small groups of Leydig cells were seen in the interstitial space (Fig.2 A, B). In LCTs, most cells possessed a large polygonal shape with abundant cytoplasm, indistinct cell borders, and regular round to oval nuclei. The nucleus was found to be frequently prominent (Fig. 2b). Occasionally, cells as those noted above were found to possess distinct cell borders and smaller nuclei (Fig. 2b’). Small cells with scant, densely eosinophilic cytoplasm and a grooved nuclei (Fig. 2b”) and spindle-shaped (sarcomatoid) cells (Fig. 2b”’) were observed as well.

**Figure 2.**
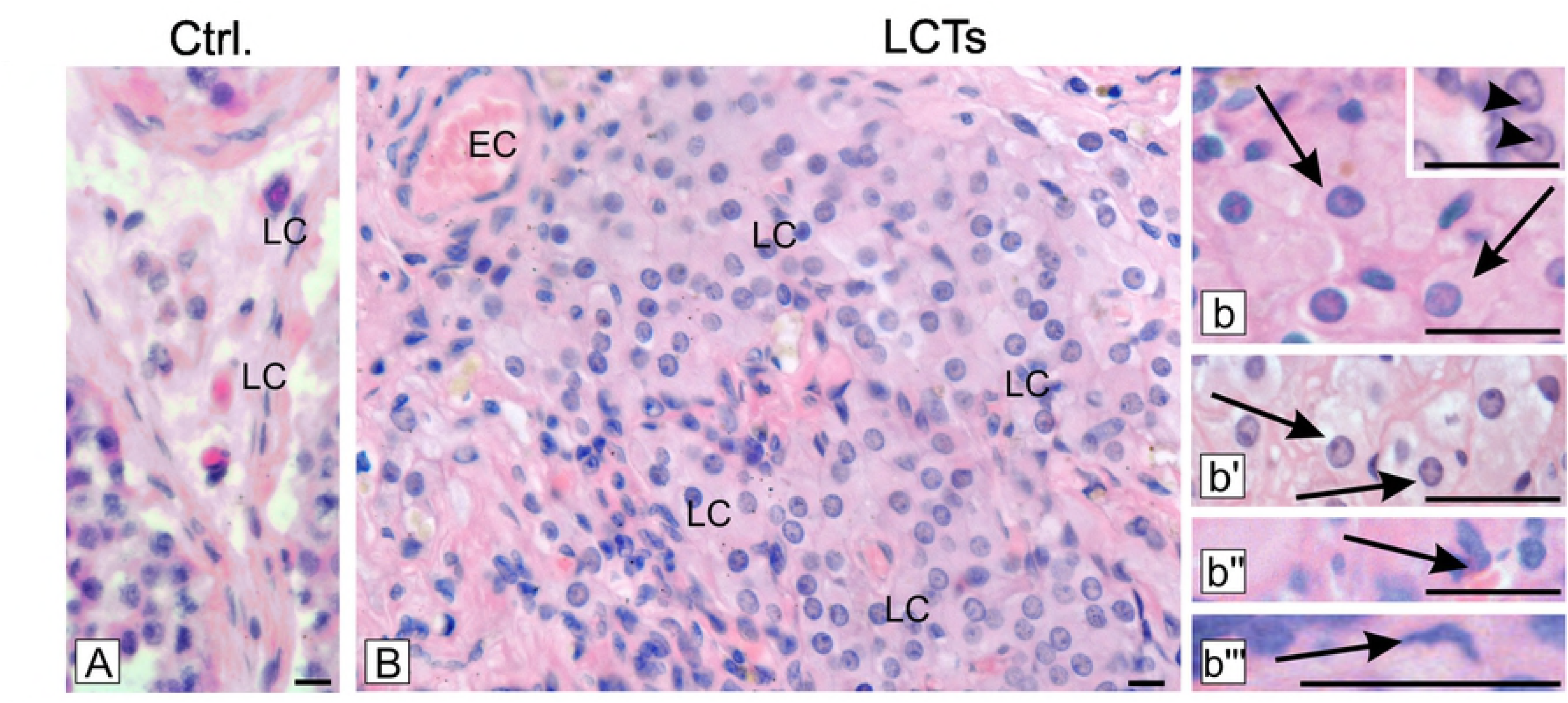
Morphology of human Leydig cell tumors – hematoxylin-eosin staining. Representative microphotographs of (A) control human testis and (B, b-b’’’) Leydig cell tumors (LCTs). Scale bars represent 15 μm. Staining was performed on serial testicular sections from n = 12 specimens. LC-Leydig cells, EC - epithelial cells of blood vessels (b) - arrows depict cells of large polygonal shape with abundant, cytoplasm, indistinct cell borders, and regular, round to oval nuclei. Prominent nucleus visible at (b) higher magnification (arrowheads), (b’) - arrows depict cells with above features but possessing distinct cell borders and smaller nuclei, (b’’) - arrows depict small cells with scant, densely eosinophilic cytoplasm and grooved nuclei, (b’’’) – arrows depict spindle-shaped (sarcomatoid) cells.

### 3.2. Expression and localization of GPER and PPARs in LCTs

In LCTs, increased expression of GPER (*p* < 0.05) and decreased expression of PPARα (*p* < 0.001), PPARβ (*p* < 0.01), and PPARγ (*p* < 0.001) was seen when compared to controls (Fig.3A, B). Corresponding to GPER and PPARs protein expression changes their mRNA expressions in LCTs are presented as supplementary material (Fig.3’).

**Figure 3.**
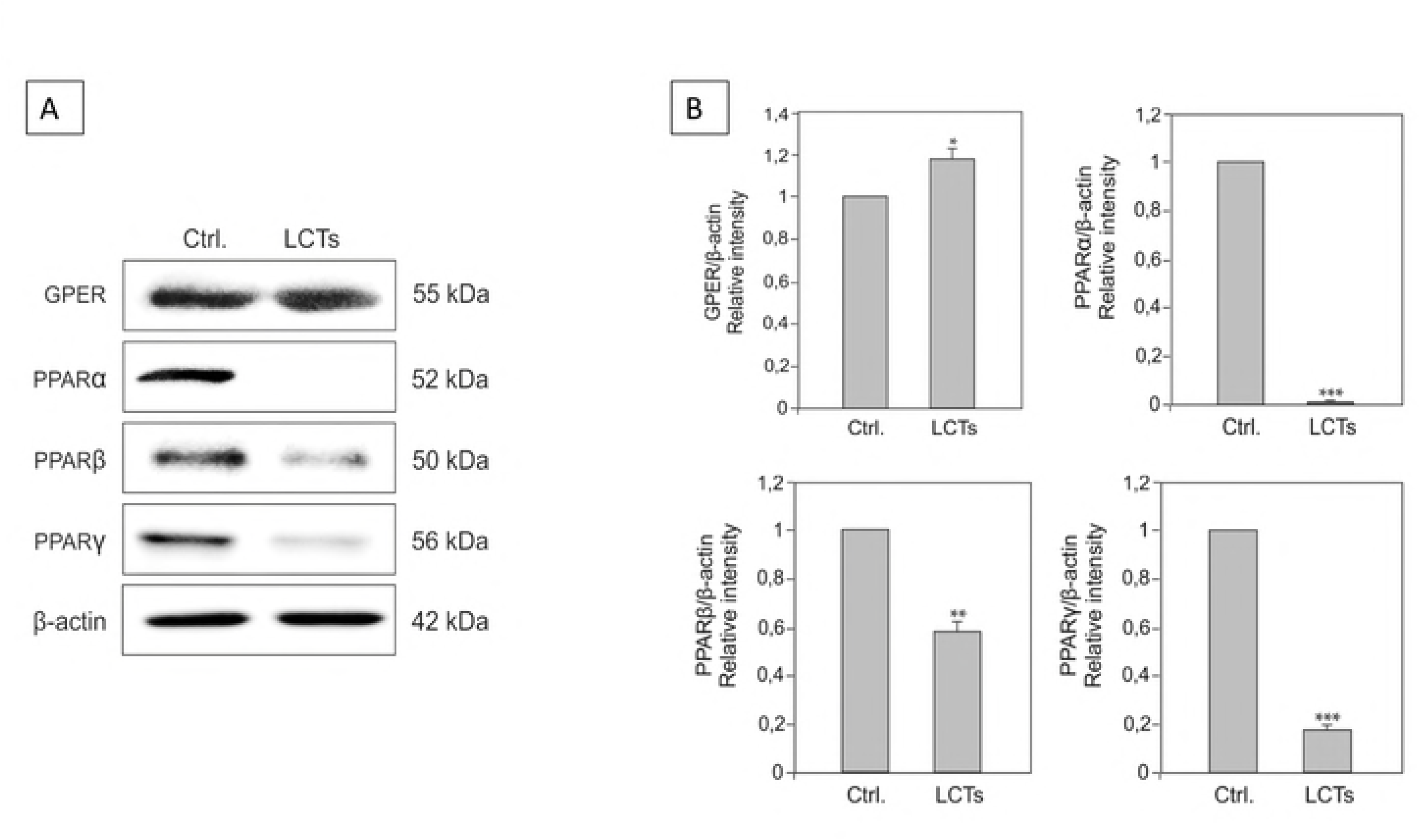
Expression of GPER, PPARα, PPARβ and PPARγ in human Leydig cell tumor. (A) Representative blots of qualitative expression of GPER, PPARα, and PPARγ and (B) relative expression (relative quantification of protein density (ROD); arbitrary units). The relative amount of respective proteins normalized to β-actin. ROD from three separate analyses is expressed as means ± SD. Asterisks show significant differences between respective control and Leydig cell tumor (LCTs). Values are denoted as ∗ *p* < 0.05, ∗∗ *p* < 0.01 and ∗∗∗*p* < 0.001. Analysis was performed in triplicate (n = 7).

**Figure 3.**
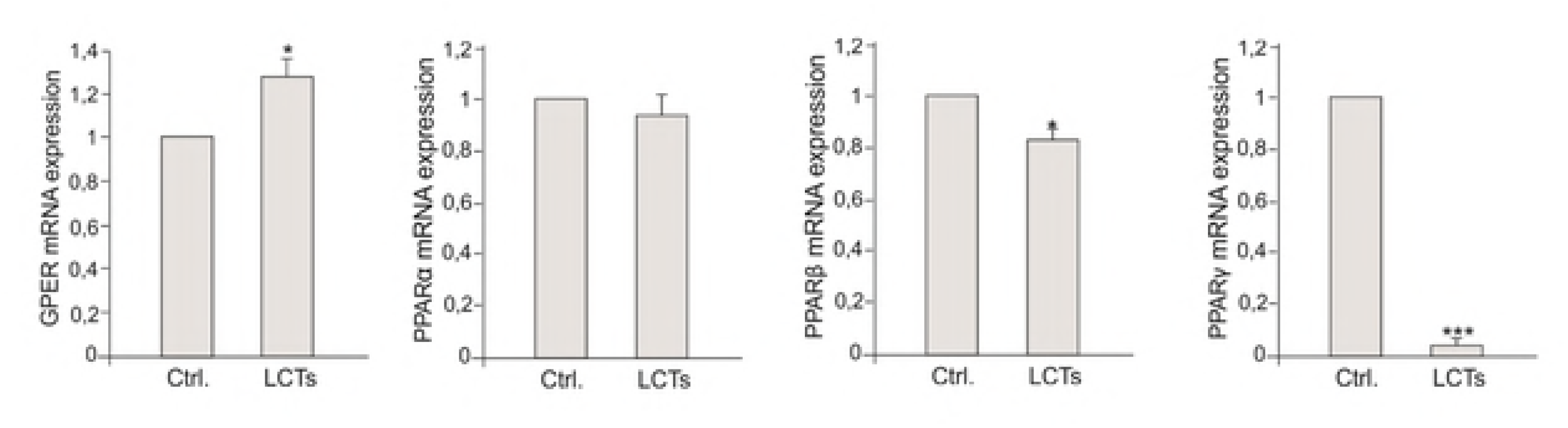
Expression of GPER, PPARα, PPARβ and PPARγ mRNA in human Leydig cell tumor. Relative level (relative quantification; RQ) of mRNA for **GPER, PPARα, PPARβ and PPARγ** determined using real-time RT-PCR analysis 2−ΔCt method. As an intrinsic control, β-actin mRNA level was measured in the samples. RQ from three separate analyses is expressed as means ± SD. Asterisks show significant differences between respective control and Leydig cell tumor (LCTs). Values are denoted as ∗ *p* < 0.05, and ∗∗∗*p* < 0.001. Analysis was performed in triplicate (n = 7).

No changes in GPER localization and staining intensity was found in control Leydig cells and LCTs (Fig.4 A, A’). Specifically, the staining was exclusively cytoplasmic and of moderate intensity. Localization and immunostaining intensity of PPAR varied between Leydig cells of control testis and LCTs (Fig.4B, B’, C, C’, D, D’). While strong cytoplasmic-nuclear expression of PPARα, β, and γ was found in control samples, an absence of PPARα expression and moderate to weak immunostaining expression of PPARβ and PPARγ, respectively, were detected. In LCTs, PPARs were located primarily in the cytoplasm of Leydig cells. No positive staining was found when primary antibodies were omitted (Fig. 4, inserts at A, D’).

**Figure 4.**
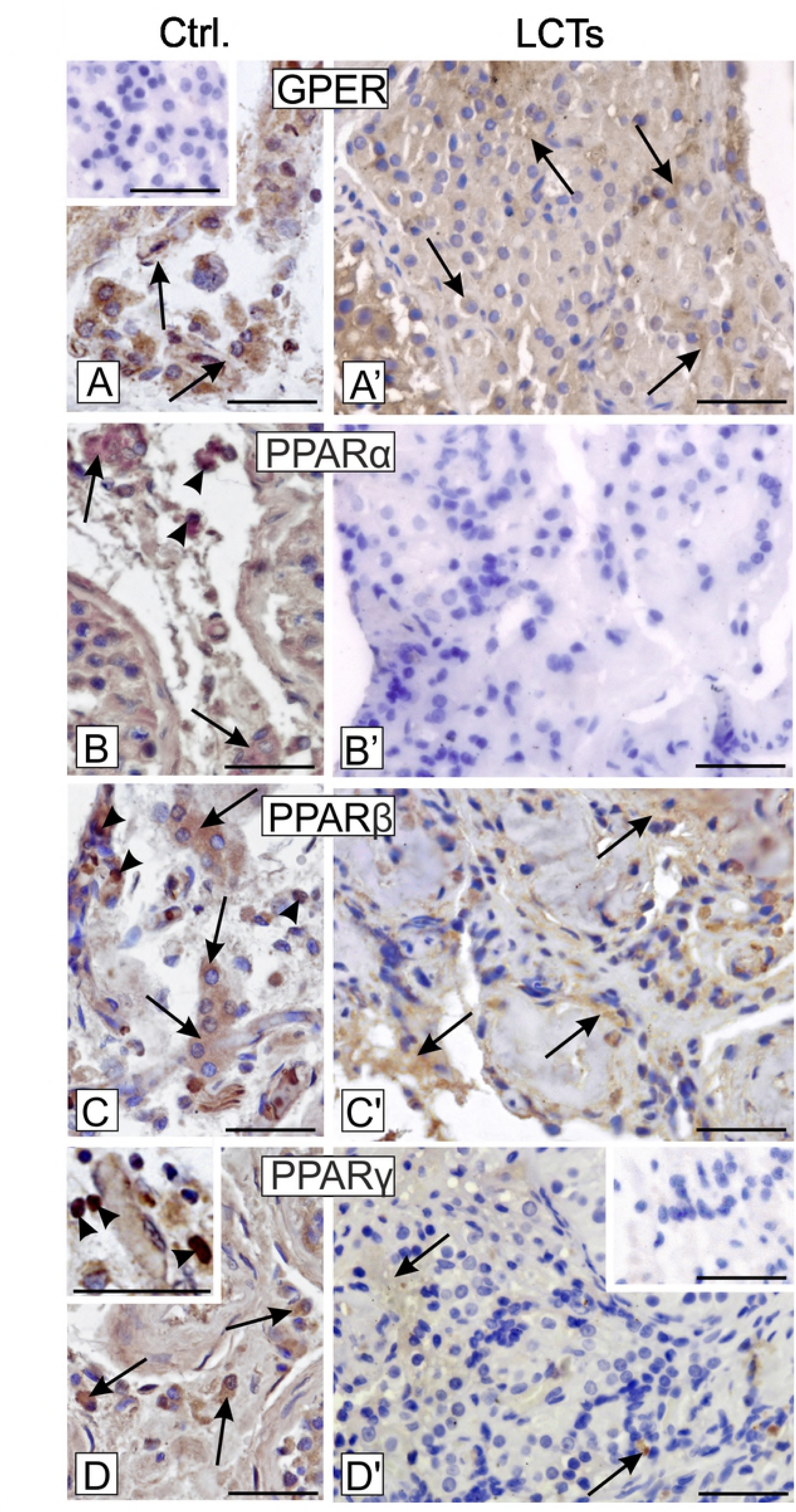
Localization of GPER, PPARα, PPARβ and PPARγ in human Leydig cell tumor. Representative microphotographs of cellular localization of GPER (A, A’), PPARα (B, B’), PPARβ (C, C’) and PPARγ (D, D’ and higher magnification at D) in control human testes (A-D and higher magnification at D) and Leydig cell tumor (LCTs). DAB immunostaining with hematoxylin counterstaining. Scale bars represent 15 μm. Staining was performed on serial testicular sections from n = 12 specimens. Arrows depict cytoplasmic staining, arrowheads depict nuclear staining. No positive staining is seen when the primary antibodies were omitted – insert at A and D’-(negative controls).

### 3.3. Expression and localization of LHR, PKA, PLIN, HSL, StAR, TSPO, HMGCS and HMGCR in LCTs

In LCTs, varying expression of LHR, PKA, PLIN, HSL, StAR, TSPO, HMGCS, and HMGCR was observed when compared to normal Leydig cells (Fig. 5A, B). The expression of LHR and PKA was increased (*p* < 0.05 and *p* < 0.01, respectively) as well as that of HMGCS and HMGCR (*p* < 0.001 and *p* < 0.05, respectively). In contrast, PLIN and StAR expression was decreased (*p* < 0.001 and *p* < 0.05, respectively), while a non-significant increase was observed for HSL and TSPO. Corresponding to LHR, PKA, PLIN, HSL, StAR, TSPO, HMGCS, and HMGCR protein expression changes their mRNA expressions in LCTs are presented as supplementary material (Fig.5’).

**Figure 5.**
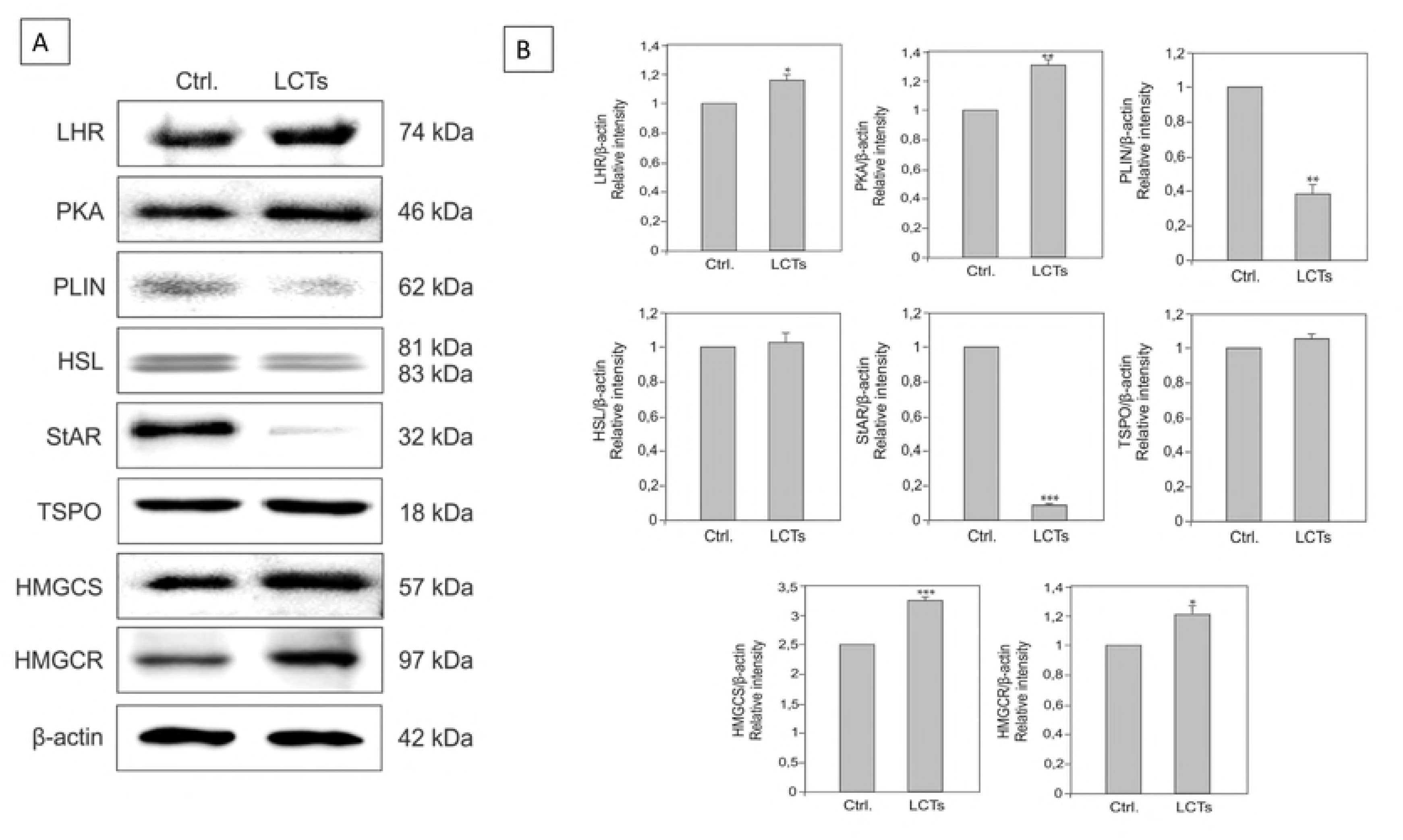
Expression of LHR, PKA, PLIN, HSL, StAR, TSPO, HMGCS and HMGCR in human Leydig cell tumor. (A) Representative blots of qualitative expression of LHR, PKA, PLIN, HSL, PLIN, StAR, TSPO, HMGCS and HMGCR and (B) relative expression (relative quantification of protein density (ROD); arbitrary units). The relative amount of respective proteins normalized to β-actin. ROD from three separate analyses is expressed as means ± SD. Asterisks show significant differences between respective control and Leydig cell tumor (LCTs). Values are denoted as ∗ *p* < 0.05, ∗∗ *p* < 0.01 and ∗∗∗*p* < 0.001. Analysis was performed in triplicate (n = 7).

**Figure 5.**
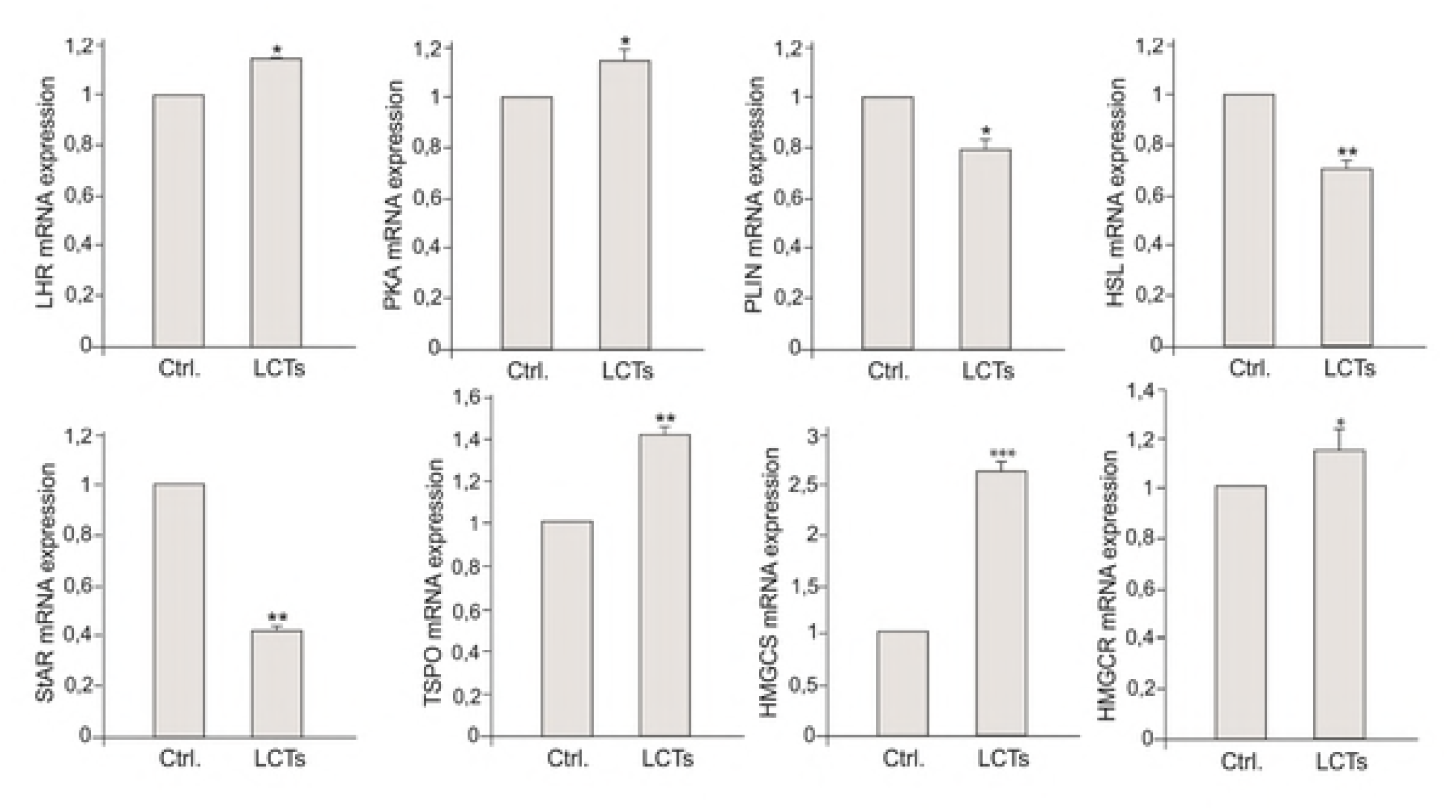
Expression of LHR, PKA, PLIN, HSL, PLIN, StAR, TSPO, HMGCS and HMGCR mRNA in human Leydig cell tumor. Relative level (relative quantification; RQ) of mRNA for LHR, PKA, PLIN, HSL, PLIN, StAR, TSPO, HMGCS and HMGCR determined using real-time RT-PCR analysis 2−ΔCt method. As an intrinsic control, β-actin mRNA level was measured in the samples. RQ from three separate analyses is expressed as means ± SD. Asterisks show significant differences between respective control and Leydig cell tumor (LCTs). Values are denoted as ∗ *p* < 0.05, ∗∗ *p* < 0.01 and ∗∗∗*p* < 0.001. Analysis was performed in triplicate (n = 7).

In control Leydig cells and LCTs, cytoplasmic expression of LHR was found (Fig.6 A, A’). The immunostaining was of moderate intensity in control Leydig cells but was weak and present in minority of cells of LCTs. No differences were found in PKA distribution and immunostaining (Fig. 6 B, B’), with strong staining present in control and tumor Leydig cell cytoplasm. PLIN distribution was cytoplasmic in control Leydig cells and LCTs (Fig. 6 C, C’). In control Leydig cells, staining intensity was strong while found to be weak in LCTs. Increased HSL staining intensity was found in LCTs when compared to control cells (Fig. 6D, D’) and was exclusively cytoplasmic. Strong immunoreaction was found in the blood vessel epithelium. In contrast, decreased staining intensity of StAR, exclusively present in the cytoplasm, was observed in LCTs (Fig. 6E, E’) while control Leydig cells exhibited moderate cytoplasmic staining. A similar pattern was found for TSPO (Fig. 6 F, F’). Moderate cytoplasmic expression was revealed in control Leydig cell cytoplasm, while the TSPO staining intensity was very weak in LCTs. However, in a few cells immunoreaction was very strong. No differences were found in the distribution of HMGCS and HMGCR between control cells and LCTs (Fig. 6 G, G’ and H, H’). Strong cytoplasmic expression of HMGCS and moderate cytoplasmic expression of HMGCR were revealed in control Leydig cells and LCTs, respectively. No positive staining was found when primary antibodies were omitted (Fig. 4, inserts at A, F’).

**Figure 6.**
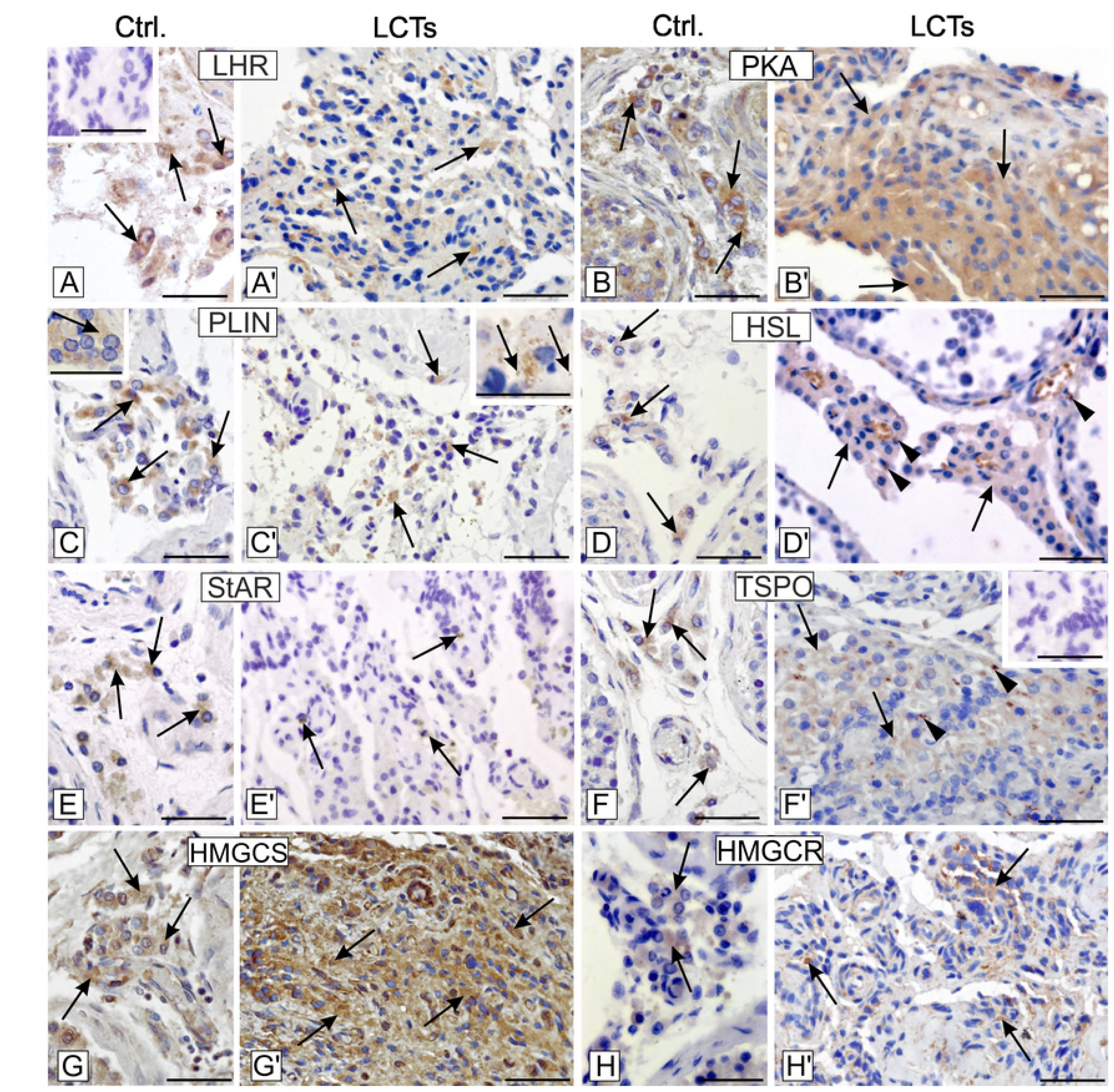
Localization of LHR, PKA, PLIN, HSL, StAR, TSPO, HMGCS and HMGCR in human Leydig cell tumor. Representative microphotographs of cellular localization of LHR (A-A’), PKA (B-B’), PLIN(C-C’ and higher magnifications at C and C’), HSL (D-D’), StAR (E-E’), TSPO (F-F’), HMGCS (G-G’) and HMGCR (H-H’) in control human testes (A-H and higher magnification at C) and Leydig cell tumor (A’-H’ and higher magnification at C’). DAB immunostaining with hematoxylin counterstaining. Scale bars represent 15 μm. Staining was performed on serial testicular sections from n = 12 specimens. Arrows depict cytoplasmic staining. Arrowheads depict strong stained cells for TSPO and positively stained epithelial cells of blood vessels for HSL. No positive staining is seen when the primary antibodies were omitted – insert at A and F’-(negative controls).

### 3.4. Effect of GPER and PPAR blockage on expression of PI3K, Akt and mTOR in LCTs

In LCTs, PI3K and Akt expression was increased (*p* < 0.05,) while no observable changes in mTOR expression was found when compared to controls (Fig. 5A, B).

### 3.5. Effect of GPER and PPAR blockage on cholesterol concentration, estradiol secretion and cGMP concentration in MA-10 cells

Regardless of used antagonists (alone or in combination), an increased (*p* < 0.05, *p* < 0.01) cholesterol concentration in tumor Leydig cells was seen (Fig. 8A). Secretion of estradiol markedly increased (*p* < 0.001) after GPER blockage (Fig.8B). A similar increase (*p* < 0.01) was observed after GPER and PPARα blockage. Conversely, blockage of GPER and PPARγ decreased (*p* < 0.05) estradiol secretion. When either PPARα or PPARγ was blocked, no to little alterations (*p* < 0.05) in hormone secretion were revealed.

Changes in cGMP concentration after antagonist-treatment were similar to those of estradiol secretion (Fig.8C). Treatment with a GPER antagonist, alone or in combination with a PPARα antagonist, increased (*p* < 0.05, *p* < 0.01) cGMP concentration while treatment with PPARα or γ antagonists consistently decreased (*p* < 0.05) the concentration. Only treatment with GPER and PPARγ antagonists in combination increased (*p* < 0.01) cGMP concentration.

## Discussion

In the present study, we examined the cellular organization and molecular mechanisms, including GPER and PPAR signaling, lipid balance, and steroidogenesis-regulating molecular interactions that regulate LCT biology.

For the first time, scanning electron microscopic analysis was used to visualize the general organization of LCTs. We showed a complicated LCT structure where individual cells were not recognized in the solid mass, but a number of prolongations of various size were formed, keeping cells tightly linked to each other. Of note, according to Kim *et al.,* [14] and Al-Agha and Axiotis [56], benign LCTs classically present as a small (3–5 cm in diameter), sharply delineated and solid mass embedded within the testis. Alternatively, malignant LCTs are typically larger (greater than 5 cm in diameter), have infiltrative margins, and show areas of hemorrhage and necrosis. They replace the testis and/or extend beyond testicular parenchyma. Morphologically, LCTs can consist predominantly of one type or as a mixture of the four types of cells [57] observed in specimens examined herein. Pale to clear cytoplasm of LCTs, related to abundant lipid accumulation, was reported by Al-Agha and Axiotis [56]. Other characteristic frequently observed is a rich cytoplasmic lipofuscin pigment that is distinctive for LCT, although it is present in other steroid hormones-secreting tumors as well as in aging cells [3]. Additionally, Reinke crystals, both intracytoplasmic and intranuclear, are often described in LCTs. Ultrastructural studies of LCTs revealed organelle content typical for steroid-secreting cells e.g. abundant smooth endoplasmic reticulum, mitochondria with tubulovesicular cristae, and numerous lipid droplets or irregularly shaped electron-dense bodies consisting of lipid droplets and accumulated in lysosomes [58].

Leydig cell tumors are sex steroid hormone-secreting tumors with androgens produced in prepubertal children and estrogens in adults [10]. A central factor in LCT growth and progression is represented by an inadequate intratesticular balance in the androgen/estrogen ratio [59]. Sirianni *et al.,* [60] demonstrated that estrogens elicit proliferative effects in human and rodent tumor Leydig cells through an autocrine mechanism. Findings from transgenic mice revealed an increased estrogen/androgen ratio, and estrogen excess resulted in Leydig cell hyperplasia, hypertrophy, and adenomas [61]. Of note, these effects can be induced by the action of estrogen *via* canonical estrogen receptors. Varying expression patterns of ERα and ERβ were observed in human LCTs compared to healthy testis [36]. Also, in human and rat LCTs, involvement of GPER in cell proliferation, growth, and apoptosis was shown [62]. Rago *et al.,* [63, 64] confirmed the presence of GPER in germ cell tumors and sex-cord stromal tumors. From the latter group, in LCTs, the authors found no differences in GPER expression in relation to normal testis. In contrast, we revealed an increase in GPER expression in LCTs when compared to normal Leydig cells. Moreover, in preliminary *in vitro* experiments in mouse tumor Leydig cells, GPER expression was increased [54].

Herein, we showed GPER, alone and together with PPARα, effected estradiol secretion by tumor Leydig cells, although GPER and PPAR increased cholesterol levels in these cells. Such results indicate a leading role of GPER over PPAR in regulation of sex hormone production and secretion, and it suggests possible GPER and PPARα alterations in LCTs. Similarly, our prior study also showed progesterone secretion modulation in GPER and PPAR antagonists-treated tumor mouse Leydig cells [52]. According to findings by Chimento *et al.,* [62], GPER is a good target for reduction of tumor Leydig cell proliferation. In tumor rat Leydig cells (R2C), GPER activation by its agonist (G-1) was associated with the initiation of the intrinsic apoptotic mechanism. Other experimental studies showed that endocrine disrupting chemicals, acting through GPER signaling, are involved in testicular germ cell carcinogenesis [37].

Interestingly, androgens have been shown to inhibit tumor Leydig cell proliferation by opposition to self-sufficient *in situ* estrogen production in R2C cells [46]. Androgen treatment significantly decreased the expression and activity of estrogen synthase (aromatase). This inhibitory effect relied on androgen receptor (AR) activation and involved negative regulation of the aromatase gene (*CYP19*) transcriptional activity through the nuclear orphan receptor DAX-1 (dosage-sensitive sex reversal, adrenal hypoplasia critical region, on chromosome X, gene 1). Alternatively, the negative role of anabolic androgenic steroids in supraphysiologic dosage was recently implicated in mechanisms of tumorigenesis *via* impairment of the expression of steroidogenic enzymes and effects on intratesticular hormonal balance [65]. Of note, male patients with congenital adrenal hyperplasia (CAH) can develop bilateral testicular adrenal rest tumors (TARTs). These tumors, in most cases, regress with glucocorticoid therapy but histological differentiation from Leydig-cell tumors is quite difficult [66]. Ulbright *et al.,* [3] reported unusual features of LCTs e.g. adipose differentiation, calcification with ossification, and spindle-shaped tumor cells in both young and ageing patients. Awareness of these features may prevent misinterpretation.

Two standards are currently used to distinguish between hyperplastic nodules from adenomas. An adenoma classification is warranted if its diameter exceeds either one seminiferous tubule cross-section [67] or three seminiferous tubule cross-sections [69]. Leydig cell hyperplasia (focal or diffuse) and adenomas are commonly observed in laboratory rodents. The spontaneous incidence of adenomas in ageing Sprague-Dawley and inbred Fischer 344 rats, as well as mature CD-l and B6C3Fl mice, have been documented [68]. Morphologically, no differences appear between spontaneous or chemically induced Leydig cell adenomas [69].

Increased PPAR expression in organ pathophysiology e.g. liver, heart, intestine, and renal proximal tubules, is currently only partially elucidated [70-73]. We showed, for the first time, a PPAR expression pattern in normal human Leydig cells and its prominent down regulation in LCT. Immunoreactive PPARα and β were clearly apparent in testicular germ cell tumors [74]. A similar correlation was found in dog testis, and PPAR expression was always markedly higher in tumor tissue [75]. Recently, Kadivr *et al.,* [76] reported a relationship between PPAR mRNA expression and spermatozoa motility in rams. Notably, confusing results were seen concerning the involvement of PPAR in tumor biology. PPAR was revealed to both promote and inhibit cancer *via* effects on cell differentiation, growth, metastasis, and lipid metabolism [77].

In our findings, both *in vivo* and *in vitro* studies revealed a partnership between GPER, PPAR, and lipid homeostasis-controlling molecules in LCT. These molecules showed altered expression patterns in relation to GPER and PPAR expression in LCT.

The effect of either GPER or PPAR on cholesterol content suggests alterations in cholesterol synthesis and/or storage that may be based on GPER and/or PPAR disturbances. Recent studies have also linked lipids abundance with increased tumor aggressiveness and its resistance to chemotherapy [78]. Wang *et al.,* [79] reported high cholesterol content and infertility in HSL knockout mice; however, no information on Leydig cell function was provided. Our studies revealed HSL expression was not disturbed in LCTs. Findings by Christian *et al.,* [80] showed that autophagy influences lipid metabolism *via* both lipogenesis (supporting cell growth within nutrient-limited areas, thereby contributing to tumor symbiotic relationships) and lipolysis. Lipid droplets may induce autophagy and undergo lipophagy to avoid lipotoxicity, a phenomenon caused by excessive lipid accumulation with involvement of the mTOR signaling pathway [81]. Thus, we show increased cholesterol content without activated mTOR can indirectly result in lipophagy induction in LCTs. It seems this particular tumor can have a distinct biology, but it is possible that some mechanisms can be induced later when its development is more advanced.

Lastly, findings by Ma *et al*., [82] demonstrated that culture of rat Leydig cells in hypoxic conditions decreased cholesterol content. These findings indicate that, besides well-known lipid homeostasis controlling molecules, a number of factors of various origin/nature are implicated in its regulation. In steroidogenic cells, the mechanism underlying lipid turnover and receptor involvement remained unanswered [77] however, based on these results, GPER-PPAR cross-talk should be taken into consideration. The question also arises whether some of these factors and/or molecules, when disturbed, lead to Leydig cell tumorigenesis with additional perturbations in lipid homeostasis/steroidogenesis, or whether perturbations in lipid homeostasis-controlling factors/molecules occur as a result of tumor initiation and development. Based on all of the aforementioned results, both mechanisms are equally possible.

The color of LCTs usually ranges from brown to yellow to gray-white depending on the lipid content of the tumor [83]. The golden-brown appearance (usually imparted by the abundant lipofuscin pigment of tumor cells) is very characteristic. According to our results, in LCTs, lipid homeostasis is disturbed as seen for other endocrine and non-endocrine tumors [84]. Various changes were revealed in expression and localization of lipid balance-controlling molecules, including those controlling steroidogenesis as well as phosphatidylinositol-3-kinase (PI3K)-Akt-mTOR signaling pathway. Scarce data concerning lipid homeostasis and/or its controlling molecules in rodent tumor Leydig cells are available.

The steroidogenic function of Leydig cells can be modulated in various physiological conditions. *In vivo* studies in young and ageing rats revealed increased sex steroid synthesis in animals administered with a TSPO ligand (FGIN-1-27) [85]. In both age groups, serum testosterone levels increased significantly. Herein, HSL and TSPO expression did not vary in LCTs, suggesting a subordinate role of these molecules in LCTs. In the Leydig cell line (M5480), an acute effect of hCG was observed as increased metabolism of cholesteryl esters was reported [87]. Moreover, in patients with testicular cancer, hCG treatment caused excess of estradiol secretion by the tumor [88]. In a mouse tumor Leydig cell line (mLTC-1), epidermal growth factor increased StAR activity and steroid production efficiency in a time- and dose-dependent manner with involvement of ERK, while LHR expression was significantly reduced [88]. In mice (C57BL/6J) with LCTs, cessation of steroidogenesis was present when LH and cAMP were removed [89]. We found prominent changes in LHR and PKA expression, reflecting disturbances in lipid controlling mechanisms directly associated with central endocrine regulation and the local microenvironment. For example, due to implications of HMGCR in cancer cell proliferation and cooperation with Ras signaling, HMGCR is used as a molecular target of statins, cholesterol-lowering drugs [90].

Metabolic flux and availability of lipids is controlled directly by lipid droplets and peroxisomes. These lipids serve as membrane stabilizers and structural elements, protein modifiers and signaling molecules, as well as energy sources for cell growth, migration, and invasion [91,92]. Therefore, failure of one of the lipid machinery components results in a negative effect on global cell physiology and lipid homeostasis. In pancreatic cancer, the content of lipid droplets is mobilized under a restricted cholesterol-rich, low-density lipoprotein (LDL) supply. Limiting LDL uptake reduces the oncogenic properties of this cancer and renders it more sensitive to cytotoxic drugs [93].

It is worth noting that lipid droplet associated proteins are actively involved in modulating lipid homeostasis by generating sites for steroidogenic enzyme activity [94, 95]. Marked expression changes in PLIN reflect that it can affect steroidogenic enzymes, thereby aiding in the development of LCTs. Upon stimulation of mLTC-1 cells with LH or 8-bromo-cAMP, large lipid droplets become much smaller and are dispersed throughout the cytosol. Lukyaenko *et al*., [95] demonstrated lipophilic macrophage-derived factor as a highly active, acute regulator of steroidogenesis, acting through a high capacity StAR-independent pathway.

Alterations in the mitochondrial status affect cell lipid homeostasis and steroid biosynthesis efficiency [96, 97]. Mitophagy serves as an indispensable mechanism to transfer damaged mitochondria for lysosomal degradation. In LCTs, either mitochondrial function and/or mitophagy can be altered, thus affecting lipid homoeostasis as indirectly shown through perturbations in StAR expression patterns.

It is worth adding that lipids have been recognized as a component of metabolic reprogramming in tumor cells [98]. Many tumors show a reactivation of *de novo* fatty-acid synthesis for generation of membrane structural lipids; thus, they do not rely on lipids from the bloodstream [99]. Modulated lipid synthesis may also have a non-cell-autonomous role in cancer development. The growth and metastasis promotion of LCT cannot be excluded with participation of adipocytes [100]. In addition to their structural and signal transduction roles, lipids can also be broken down into bioactive lipid mediators, regulating a variety of carcinogenic processes including cell growth, cell migration, and metastasis formation, as well as the uptake of chemotherapeutic drugs [101, 102]. Therefore, based on our results that show alterations in various lipid-controlling molecules, further studies are needed to elucidate the type, role, and regulation of lipids synthesized in tumors of steroidogenic cells. In prostate cancer progression, stimulatory effect of insulin on steroidogenesis was reported [103].

Enhanced expression of sterol regulatory element binding proteins (SREBPs), involved in cholesterol and fatty acids synthesis through the Akt pathway (anchored-lipid membrane protein), correlates with tumor development, progression, and invasiveness, as well as increased lipid content in cell membranes [104]. Lipid raft disruption inhibits Akt activation thereby reducing tumor cell proliferation [105]. The tumor microenvironment has an essential role in the metabolic adaptation of cancer cells [77]. Through both the post-translational regulation and induction of transcriptional programs, the dysregulated PI3K-Akt-mTOR pathway coordinates the uptake and utilization of multiple nutrients e.g. lipids supporting the enhanced growth and proliferation of cancer cells [106]. In the testis, blockage of mTOR markedly decreased intracellular testosterone concentration [106]. In this study, we found PI3K, Akt, and mTOR signaling alterations present in LCTs. It is probable that, in LCTs, an increased phosphorylated and unphosphorylated kinases affect mTOR. The activation of mTOR is possible by other signaling pathway or can be related to advanced LCT development [107]. Our earlier studies in mouse tumor Leydig cells revealed that GPER and PPAR inhibition activate PI3K and Akt [52] but mTOR is modulated diversely; inhibits by GPER antagonist alone and together with PPARγ antagonist as well as activates by GPER with PPARα together and the latter alone (supplementary Fig.7’).

**Figure 7.**
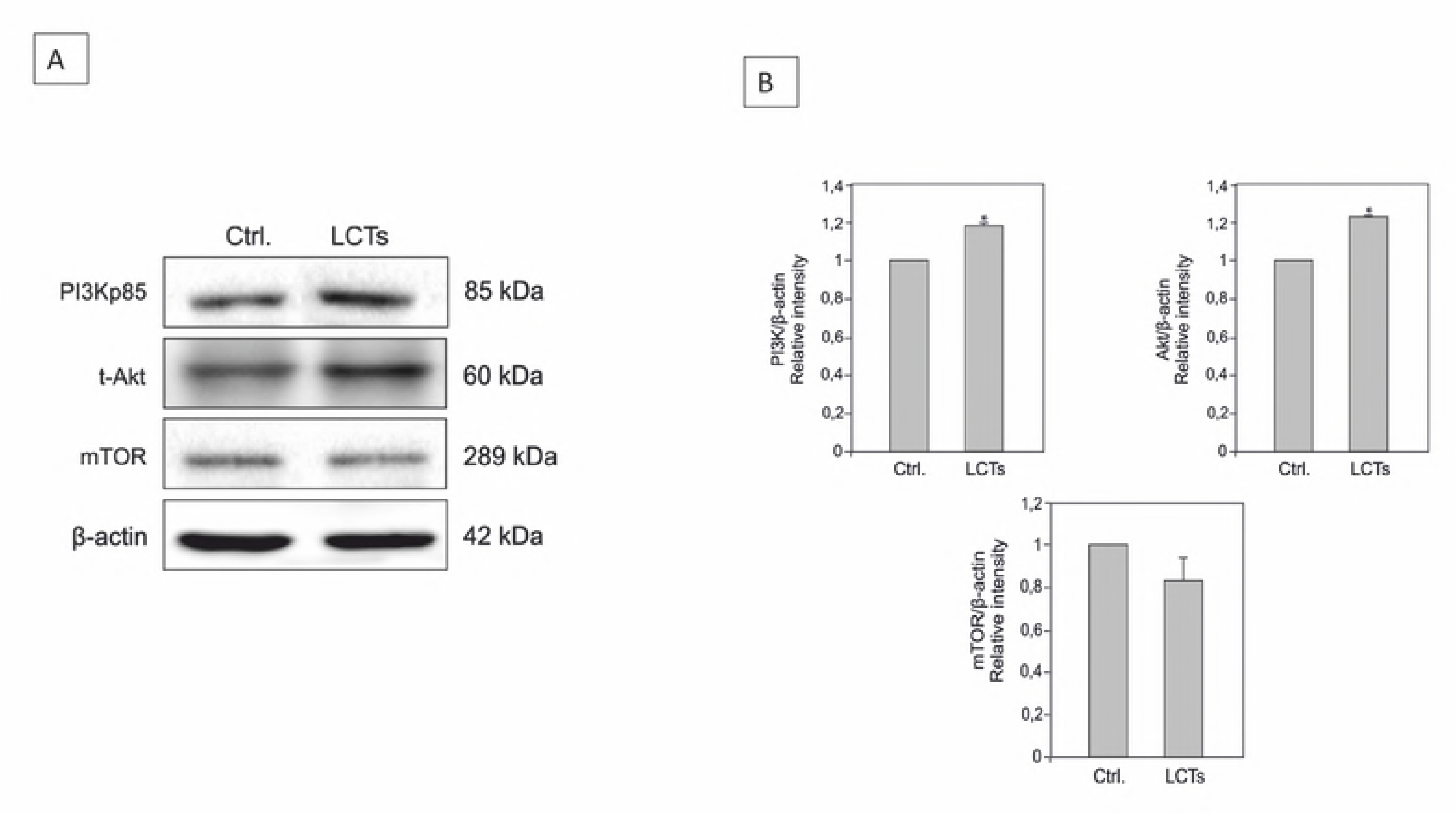
Expression of PI3K-Akt-mTOR pathway in human Leydig cell tumor. (A) Representative blots of qualitative expression of PI3K, Akt, mTOR and (B) relative expression (relative quantification of protein density (ROD); arbitrary units). The relative amount of respective proteins normalized to β-actin. ROD from three separate analyses is expressed as means ± SD. Asterisks show significant differences between control and Leydig cell tumor (LCTs). Values are denoted as ∗ *p* < 0.05. Analysis was performed in triplicate (n = 7).

**Figure 7.**
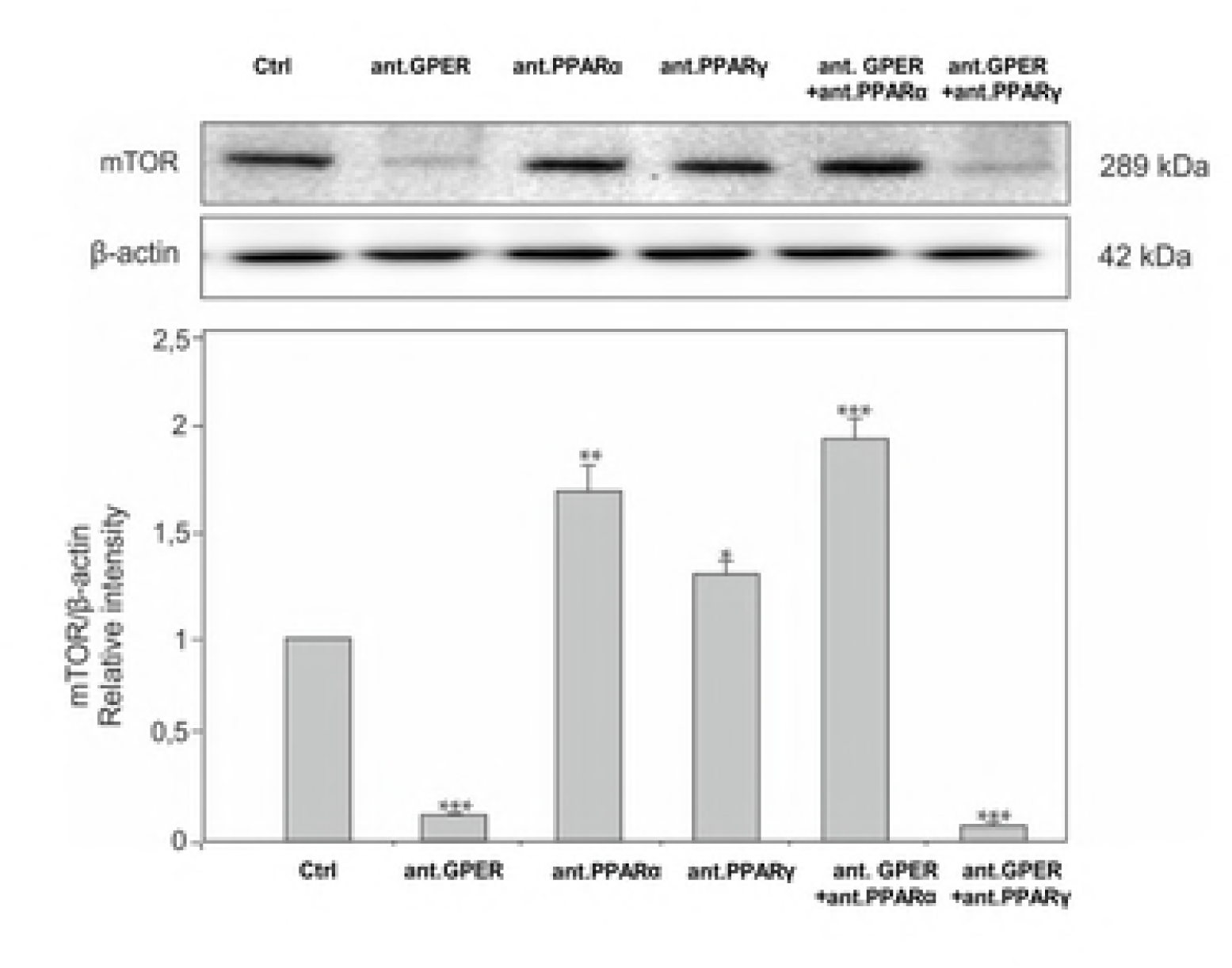
(supplementary). Effect of GPER and PPAR blockage on expression of mTOR in MA-10 cells. (A) Representative blots of qualitative expression of mTOR and (B) relative expression (relative quantification of protein density (ROD); arbitrary units). The relative amount of protein normalized to β-actin. ROD from three separate analyses is expressed as means ± SD. Asterisks show significant differences between control and treated Leydig cells. Values are denoted as ∗ *p* < 0.05, ∗∗ *p* < 0.01 and ∗∗∗*p* < 0.001. Analysis was performed in triplicate (n = 3 for each experimental group).

**Figure 8.**
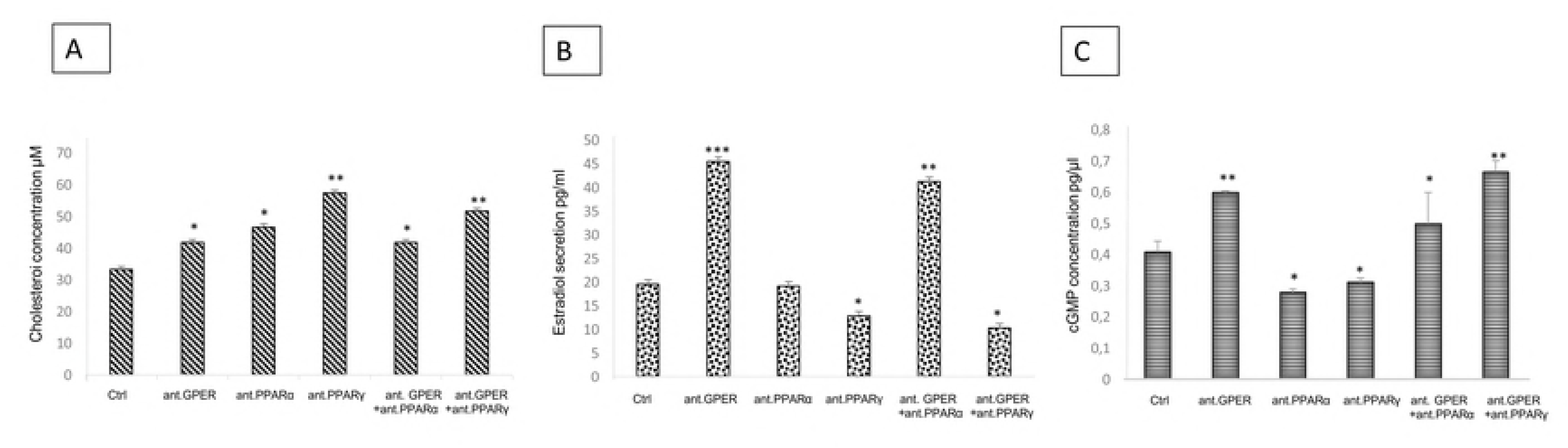
Effect of GPER and PPAR blockage on expression on cholesterol content, estradiol secretion and cGMP concentration in MA-10 cells. Cholesterol content (A), estradiol secretion (B) and cGMP concentration (C) in control and treated with GPER (10nM), PPARα (10µM) and PPARγ (µM) antagonists alone or in combinations for 24h tumor mouse Leydig cells (MA-10). Asterisks show significant differences between control and treated Leydig cells. Values are denoted as ∗ *p* < 0.05, ∗∗ *p* < 0.01 and ∗∗∗*p* < 0.001. Analysis was performed in triplicate (n = 3 for each experimental group).

Distinct changes is cGMP level suggests GPER-PPAR-mediated high or only *via* PPAR low metastatic activity of LCT. Besides antimetastatic strategy with cGMP against in colorectal cancer metastasis cGMP as a mediator of lipolysis in bovine oocytes was confirmed [108]. In LCTs, cGMP is important in maintaining lipid homeostasis when GPER and PPAR are absent.

The outcome of lipid content modification is a result of protein–protein cross-talk between messenger proteins, enzymes, and receptors that regulate pro-oncogenic and apoptotic interactions, including their activity that was revealed here also between GPER-PPAR and HMGCS and HMGCR resulting in overexpression of the latter enzymes that also is in line with our earlier study [52]. According to Ding *et al*., [109], HMGCR is an important marker for tumor testis transformation in mice.

Understanding the principles underlying these processes and mechanisms, as well as their relationship, may provide an avenue for controlling cellular lipid balance (through manipulating lipid composition), including managing post-translational modifications of proteins as well as lipid-related gene expression in LCTs [110].

## Conclusion

Mechanisms concerning Leydig cell tumorigenesis are scarce, and the role of lipid metabolism in tumor cells has long been disregarded. Recently, however, these mechanisms are recognized as the future of prominent target of therapy (Sreedhar and Zhao 2018). Therefore, alterations in lipid- and cholesterol-associated proteins and mechanisms in LCTs are presented here for the first time.

Our findings shed light on the novel functional interplay between GPER and PPAR ultimately affecting lipid metabolism and steroidogenesis in LCTs. In addition, modifications of LHR, PKA, PLIN, HSL, StAR, TSPO, HMGCS, and HMGCR, together with cGMP and PI3K-Akt-mTOR pathways, may be required in developing innovating approaches (combined with transcriptome/proteome analyses and lipidomic data) that target pathological processes of Leydig cells. There is an urgent need for additional experimental and clinical data to complete the current knowledge on the biology and molecular characteristics of LCTs. Ultimately, this will guide the early diagnostics, treatment, and surveillance of incoming patients with this disease.

## Author contributions

Authors’ contribution to the work described in the paper: M.K-B., E.G-W., M.K., P.D., A.M., P.P., W.T., B.J. P., A.H. performed research. M.K-B., E.G-W., W.T., A.H., B.B., J.K. W. analyzed the data.

M.K.-B. designed the research study and wrote the paper. All authors have read and approved the final version of the manuscript.

## Acknowledgments

The Authors are grateful to Grzegorz Kapuscinski, FEBU (urologist) and Robert Wyban, MS (LAB-IVF) (nOvum Fertility Clinic, Warszawa) for cooperation. We thank Dr. Grzegorz Wojtczak (Department of Cell Biology and Imaging, Institute of Zoology and Biomedical Research, Jagiellonian University in Kraków) for technical help in scanning electron microscopic analyses and to Miss Maja Kudrycka and Miss Patrycja Dutka (Department of Endocrinology) for technical help in qRT-PCR and Western blotting analyses.

